# Serial Dependence During Visuomotor Integration is Robust to the Passage of Time and Interference from Intervening Tasks

**DOI:** 10.1101/2025.03.17.642023

**Authors:** Esaú Ventura Pupo Sirius, André Mascioli Cravo, Raymundo Machado de Azevedo Neto

## Abstract

When intercepting a moving target, responses are systematically biased toward the time of impact from the previous trial. This phenomenon, known as serial dependence, relies on a memory mechanism that remains poorly understood. In interceptive tasks, multiple stimulus features — such as speed, time, or motor responses — can guide behavior on the current trial and may be stored to influence subsequent trials. Here, we examined how memory decays over short inter-trial intervals (Experiment 1, *N* = 23) and whether interleaved tasks influence serial dependence (Experiment 2, *N* = 28). Participants performed either a temporal reproduction task or a speed judgment task, designed to compete for temporal and speed-processing resources, respectively. Our findings reveal that serial dependence persists across all inter-trial durations and remains unaffected by intervening tasks. While serial dependence was neither reduced nor eliminated, variations in responses were partially influenced by prior temporal reproductions from the interfering task. These results suggest that serial dependence in visuomotor tasks is robust to both the passage of time and external interference, though task responses may be subtly modulated by preceding temporal reproductions.

## Introduction

Serial dependence is a behavioral bias towards judging the current stimulus similar to previously presented stimuli. This phenomenon has gained increased interest in recent years. It has been observed in several tasks and visual features such as orientation (J. Fischer & Whitney, 2014), localization (Feigin et al., 2021), color (Barbosa & Compte, 2020), numerosity (Fornaciai & Park, 2018) and faces (Liberman et al., 2014, 2018). In most of these studies, serial dependence is investigated for features actively encoded and stored in working memory in the current trial. For example, Fischer and Whitney (2014), in their seminal work, instructed participants to remember and reproduce the orientation of a Gabor patch after a retention period on every trial. They found that judgments were influenced not only by the current orientation but also by the orientations of previous trials. Critically, keeping the orientation in working memory in the current trial was an essential part of this task (J. Fischer & Whitney, 2014) and other studies (for reviews, see Manassi et al., 2023; Pascucci et al., 2023).

Although many serial dependence studies use working memory tasks, this bias has also been observed in perceptual (Fornaciai & Park, 2018; Manassi & Whitney, 2022) and visuomotor tasks (Kwon & Knill, 2013). In these experiments, participants are not required to hold information in working memory on the current trial. For example, several studies have observed serial dependence during the interception of moving targets (Bilacchi et al., 2022; De Azevedo Neto & Bartels, 2021; Kwon & Knill, 2013; Makin et al., 2008). In this kind of task, participants have to observe a target moving across the screen and press a button at the same time the target hits a barrier or crosses a hitting zone. In some studies, the collision is visible (for instance, De Azevedo Neto & Bartels, 2021); in others, it happens behind an occluder (Kwon & Knill, 2013). While responses in these studies fluctuate around the time of impact, there is a systematic error towards the time of impact in the previous trial. That is, after slower targets, responses tend to happen later in the current trial, whereas after faster targets, responses tend to happen earlier in the current trial.

In serial dependence research, a central debate is whether bias arises from perceptual or motor traces of previous trials. In visuomotor tasks, this question is complicated by the correlation between two perceptual features - target time-to-contact and speed. Specifically, faster-moving targets also have shorter times to contact when traveling the same distance. To disentangle the effects of speed, distance, and time-to-contact, Kwon and Knill (2013) randomly sampled target speeds and distances across trials to isolate the influence of each feature. Their analysis revealed that only the speed from the previous trial, rather than distance or time-to-contact, influenced responses in the current trial. However, their design conflated distance and time-to-contact, as distances were sampled to result in a range of times for each speed. They also did not assess the role of motor traces or how memory traces contributing to serial dependence might evolve between trials. Consequently, approaches that directly examine memory traces in visuomotor tasks remain limited.

In pursuit of these questions, De Azevedo and Bartels (2021) sought to distinguish perceptual from motor origins of serial dependence in a coincident timing task using Transcranial Magnetic Stimulation (TMS) to induce virtual lesions. They found that disrupting the dorsal premotor cortex between trials reduced serial biases, whereas disrupting the visual motion area (hV5/MT+) did not. This finding supports a motor origin of serial dependence; however, premotor regions have also been linked to perceptual processing of speed (Schubotz, 2007) and time (Wiener et al., 2014), leaving it unclear which specific features from previous stimuli or responses impact behavior in visuomotor tasks.

In the present study, we explored the basis of serial dependence in visuomotor integration using a different approach. Experiment 1 aimed to replicate and extend prior findings on how different inter-trial intervals (ITIs) modulate serial dependence (Bilacchi et al., 2022). Experiment 2 examined whether performing tasks interleaved with the main coincident timing task would influence serial dependence. This manipulation was based on previous research using masking (Magnussen, 2000; Magnussen & Greenlee, 1992), which suggests that serial dependence should decrease if prior trial information relies on processing resources shared with an intervening task. Participants performed either a temporal reproduction task or a speed judgment task, intended to compete for temporal and speed processing resources, respectively. A control orientation reproduction task was also included to assess whether merely interrupting participants with any task between main trials could affect serial dependence. Our results indicate that serial dependence was present across all ITI durations and persisted even with intervening tasks. Although serial dependence was not reduced or eliminated, variations in the main task responses were partially influenced by temporal reproductions from the interfering task. These findings suggest that serial dependence in visuomotor tasks is robust to the passage of time and interference, although responses themselves may be modulated by prior temporal reproductions.

## Experiment 1

### Materials and Methods

#### Participants

Twenty-three participants (12 women, 32.04±10.73 years old, mean ± standard deviation) were informed about the experimental procedures and agreed to a Consent Form provided via electronic correspondence. We recruited a convenience sample through advertising on social media. The experimental protocol was approved by The Research Ethics Committee of the Federal University of ABC. All experiments were performed following the approved guidelines and regulations. All participants had normal or corrected to normal vision and reported no acute neuropsychological disorders.

The target sample size was 24 participants, determined through a power analysis for a one-tailed t-test with 95% power and an α level of 0.005. This analysis used a Cohen’s d of 0.95, based on similar findings from the literature (Makin et al., 2008). Although three additional participants were initially recruited to account for potential attrition, the final sample size was n = 23 due to four participants not returning for the experiment’s second session.

#### Stimulus and Task

Participants performed a coincident timing task in which they were instructed to press a button at the same time a target moving from left to right at a constant speed hit a barrier (figure 1). The experiment was performed online, so screen size and room conditions varied. However, participants were instructed to be in a quiet environment and instructed via a video call by the experimenter before the first session. The experiment was built in *Psychopy* (v 2020.1.2; Peirce et al., 2019) and ran online through the *Pavlovia* platform.

**Figure 1.**
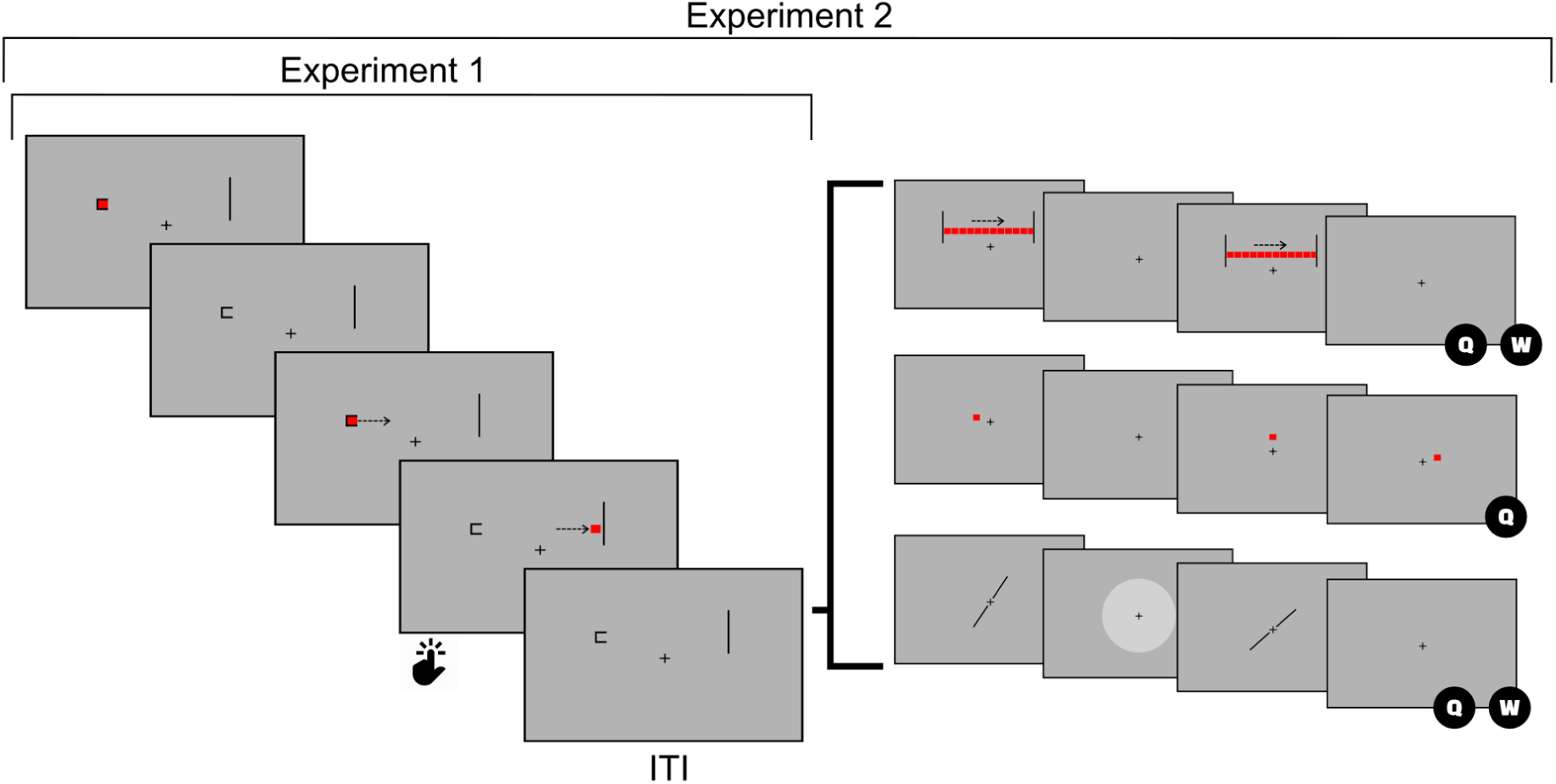
Experimental Design. *Note.* Experimental design for experiments 1 and 2, as indicated. The target from the coincident timing task would appear for 200 ms, then reappear after 200 ms, marking its trajectory’s start. Trajectories could take 496, 450, 412, 379, or 351 ms. Participants’ responses were recorded for another 500 ms after the target collision. In experiment 1, ITI was variable, taking either 0, .25, .5, 1 or 2 s. In experiment 2, ITIs was fixed at 1 s and immediately followed by inter-trial tasks depending on the experimental session.

Due to the online nature of the experiment, all sizes were defined relative to the participant’s screen height (percentage of screen height, henceforth psh). The target was a red square with a side of 3 psh and was initially drawn at the left, five psh above the vertical center of the screen. A semi-square the same size as the target marked the starting zone. The target was shown for 200 ms, marking the trial’s start, then reappeared 200 ms later, starting a trajectory towards a 30 psh vertical barrier on the right side of the screen. The target always moved at a constant speed, with interception times of 496, 450, 412, 379, or 351 ms. Interception times were defined to reflect similar times achieved in a previous experiment in the laboratory for speeds between 20 and 28 visual degrees (Bilacchi et al., 2022). The target path was set to be 60 psh and centered on the horizontal center of the screen. Participants were instructed to look at a fixation cross of the same size as the target during the trial and press the spacebar when the target hit the barrier. Responses were recorded for up to 500 ms after the target disappeared behind the barrier and were followed by an inter-trial interval (ITI).

#### Experimental Design and Procedures

Participants performed 30 mini-blocks of 51 trials each, divided into two sessions of approximately 30 minutes. Sessions were identical in the number of blocks and conditions and could be completed within a week. Within each mini-block, the target speeds were counterbalanced, such that each target speed was preceded by every other speed the same number of times (Brooks, 2012). This sequence of trials ensured that the current and previous target times to contact were independent.

Each block consisted of 5 mini-blocks of the same experimental condition. In each condition, the inter-trial intervals were fixed at 0, 0.25, 0.5, 1, or 2 s. These values were chosen to complement our previous study that looked at ITIs of 0.1, 1, 2, 4, or 8 s (Bilacchi et al., 2022). The new set of ITIs used in our current experiment had a finer grain at lower values and two overlapping ITIs with our previous experiment. Participants had to rest for at least 30 seconds between mini-blocks but were allowed longer breaks as the next mini-block started after a key press.

#### Analysis

We calculated the participant’s Temporal Error (TE) on every trial as the difference between the target time to contact and the response time in milliseconds. Negative values indicate early responses, whereas positive values indicate delayed responses. We removed outliers by calculating the median absolute deviation (MAD) for each participant and ITI condition and removing trials where the deviation was greater than 3 (Leys et al., 2013; Rousseeuw & Croux, 1993). Data from conditions where participants had more than 15% of the trials marked as bad (based on the number of trials missed and marked as outliers) were excluded from further analysis. Three of the twenty-three participants were excluded from further analysis on the 0.5 s ITI condition, one on the 1 s ITI condition, and one on all conditions.

Serial dependence was accessed by modeling the TE for each participant and ITI using multiple linear regression (equation 1), in which β_2_ is a measure of the previous trial’s influence on the current TE. We included current trial times in the regression to control for biases associated with the current trial, which is a process equivalent to the residualization commonly performed in the literature to avoid overestimating the previous trial’s influence (Bliss et al., 2017; Pascucci et al., 2019).

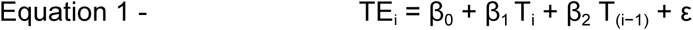

A positive β_2_ indicates an attractive serial dependence, as it means that for longer times to contact on the previous trial, participants tend to have more delayed responses in the current trial. That is, participants respond as if the current times to contact were longer than they are, therefore, more similar to the previous time to contact.

To evaluate the existence of attractive serial dependence in each ITI, a one-tailed, one-sample t-test was performed on the β_2_ coefficients. To evaluate the serial dependence dynamics between ITIs, we used a repeated measures ANOVA followed by a *post-hoc* Helmert contrast. Biases associated with the current trial were assessed using a two-tailed t-test on the β_1_ coefficients. We performed data pre-processing and regressions in Python and statistics analysis in JASP (v 0.17.2.0) and Jamovi (v 2.3.21).

In a second analysis, we combined data from our previous experiment (Bilacchi et al., 2022) with our current data. This analysis was performed to: (1) compare serial dependence on a controlled lab setting (Bilacchi et al., 2022) with our current online results; (2) have a more fine-grained set of ITIs to observe whether specific ITIs led to stronger/weaker serial dependence. We first tested if there were differences in mean and standard deviation of TEs between experiments using a two-tailed independent-samples t-test, then a similar regression was performed with current and previous contact times as predictors. To combine results from both experiments, a linear mixed model was performed with experiment type (online or laboratory) and ITIs (0, 0.1, 0.25, 0.5, 1, 2, 4, or 8 s) as categorical predictors for the obtained values of serial dependence (β_2_ coefficients) and participants were used as random variables.

### Results

Temporal errors were centered around 0 ms for all ITIs (figure 2A), indicating that participants performed the task as instructed, i.e., neither anticipating nor reacting to the target. We also plotted TEs’ means and standard deviations for each pair of previous and current times to inspect data patterns (Figure 2B and C, respectively). Figure 2B shows two general patterns: (1) participants exhibit earlier responses for longer times in the current trial, which is evidenced by the columns in Figure 2B becoming bluer vertically, and (2) participants exhibit later responses as times in the previous trial become longer, which is evidenced by cells becoming redder from left to right on every ITI. We fitted a multiple linear regression (equation 1) to evaluate these patterns separately for each participant and condition.

**Figure 2.**
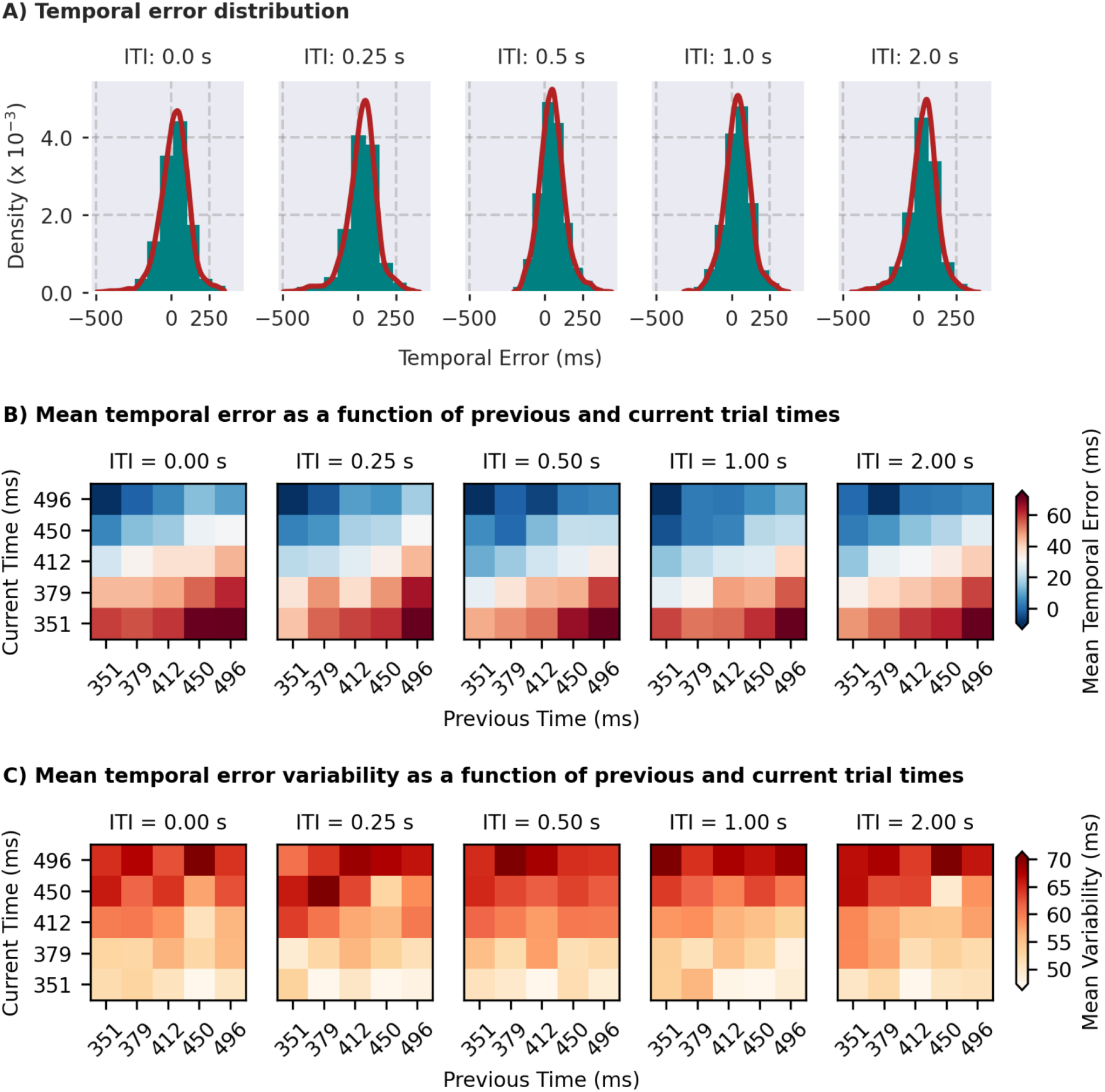
Data Overview of Experiment 1. *Note.* **A** temporal error density for all participants per inter-trial interval (ITI). The distribution around 0 and between 250 and -250 milliseconds show that participants are performing the task as expected. **B** Mean temporal error for all volunteers as a function of previous (horizontal axis) and current (vertical axis) target times. Participants tend to anticipate targets that take longer to hit the barrier (cells get bluer from bottom to top), but also tend to delay answers as the previous target time increases (cells get redder from left to right). **C** Mean temporal error variability for all volunteers as a function of previous (horizontal axis) and current (vertical axis) target times. Responses become more variable as current target times increase (cells get redder from bottom to top), consistent with the scalar property of time.

The estimated coefficients for serial dependence can be seen in figure 3A. We found statistically significant attractive serial dependencies for all ITIs, evidenced by a one-sample one-tailed t-test on the previous time coefficients [0 s: t (21) = 7.589, Cohen’s d = 1.618; 0.25 s: t (21) = 7.929, Cohen’s d = 1.690; 0.5 s: t (18) = 7.422, Cohen’s d = 1.703; 1 s: t (20) = 7.163, Cohen’s d = 1.563; 2 s: t (21) = 8.250, Cohen’s d = 1.759, all p-values <.001]. As previous times become longer, the current temporal error becomes more positive, meaning that participants respond to the current stimuli as if they were more similar to past stimuli. We also compared the magnitude of serial dependence across ITIs and, contrary to our previous results (Bilacchi et al., 2022), found no statistically significant difference [F(4,72) = 1.840, p = .130, ⍵² = 0.023]. However, repeated measures ANOVA does not account for the ordinal nature of the tested inter-trial intervals (ITIs), which may reduce its sensitivity to detect differences, especially with the higher variability typical of online data. Because of this, we performed an exploratory analysis using a Helmert contrast to compare the serial dependence in increasingly longer ITIs to the mean of all subsequent ITIs. We found a reduction in serial dependence after 0.25 s [Helmert contrast between 0.25 s and all longer ITIs: t(21) = 2.048, p = .044]. Although this result must be interpreted cautiously, given its exploratory nature, it captures a decrease in serial dependence consistent with the previously detected time window between 0.1 s and 1 s (Bilacchi et al., 2022).

**Figure 3.**
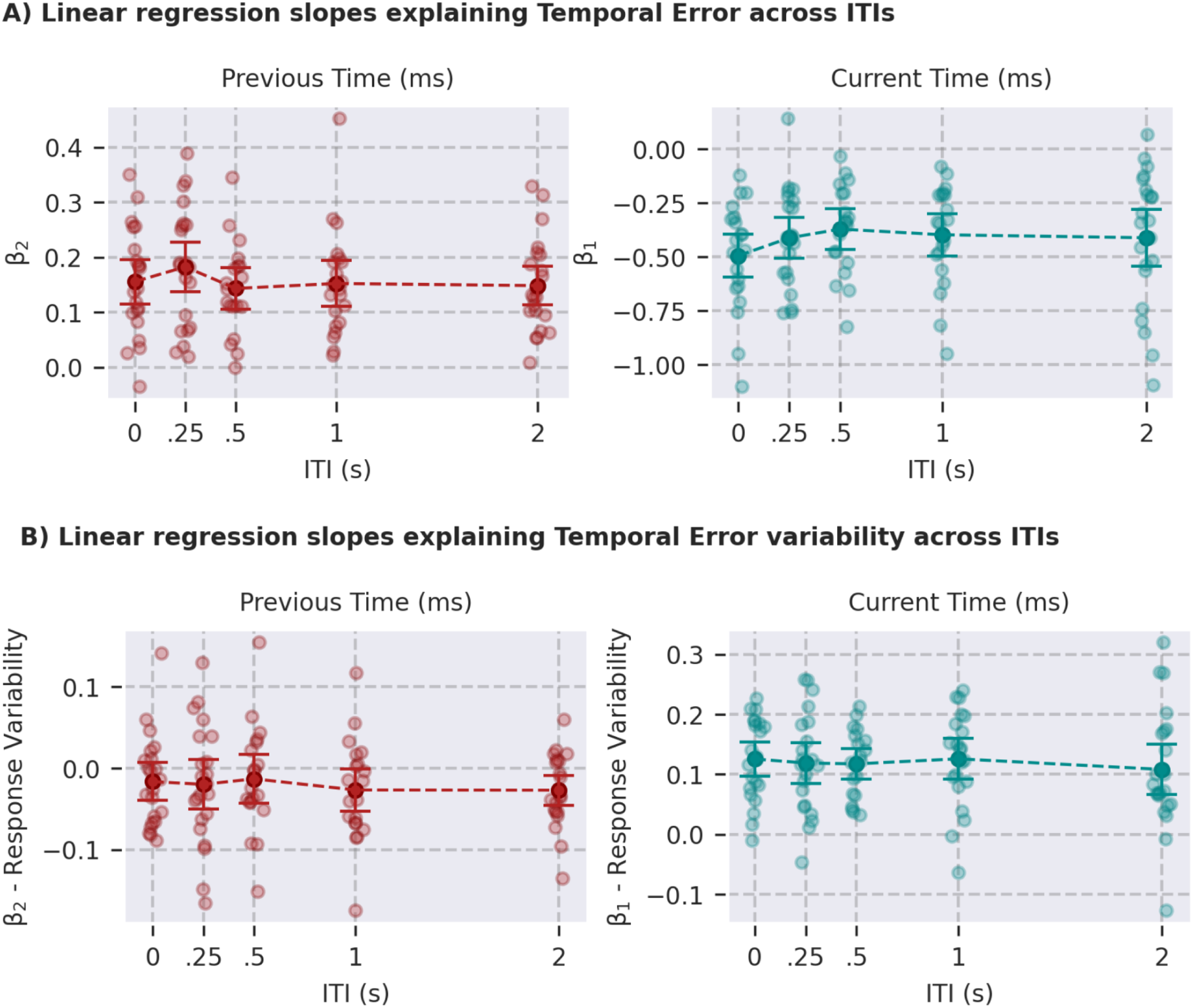
Linear Regression Slopes for Experiment 1. *Note.* Previous (left) and current (right) time slopes of multiple linear regressions performed on individual participant data for all ITIs. Large opaque dots represent average parameter estimates across participants, small and transparent dots represent individual data, and error bars represent 95% confidence intervals over parameter estimates. **A** shows slopes for the linear regression performed on temporal errors. **B** shows slopes for linear regression performed on temporal error variability, defined as the standard deviation of temporal errors for each pair of current and previous times.

Our previous study found a bias towards earlier responses for longer times to contact - or faster speeds - in the current trial. A similar result was observed in this dataset [0 s: t (21) = -9.628, Cohen’s d = 2.053; 0.25 s: t (21) = -8.512, Cohen’s d = 1.815; 0.5 s: t (18) = -7.713, Cohen’s d = 1.770; 1 s: t (20) = -7.965, Cohen’s d = 1.738; 2 s: t (21) = -6.113, Cohen’s d = 1.303, all p-values <.001], evidenced on the pattern of figure 2B. Contrary to the results of Billachi et al. (2022), a difference was found between current trial biases across ITIs [F(4,72) = 2.599, p = .043, ⍵² = 0.017] (figure 3). This was due to a larger bias detected when there was no interval between the trials [Helmert contrast between 0 and all other ITIs: t(21) = -2.818, p = .006], a condition absent in the past study.

In a second analysis, we investigated response variability as a function of ITIs. For every pair of current and previous times, per ITI, the mean standard deviation is plotted in figure 2C. There is no distinguishable pattern except for an increased variability for longer current times. There was no difference between the standard deviation of temporal errors per ITI [F(4,72) = 1.522, p-value = .205, ⍵² = 0.002]. To investigate the response variability further, we used a similar modeling approach with the multiple linear regression in equation 1, with standard deviation as the dependent variable instead of TEs (figure 3B). We found that responses become more variable as current times get longer for all ITIs, which is evidenced by β_1_ coefficients that are significantly different from zero [0 s: t (21) = 8.612, Cohen’s d = 1.836; 0.25 s: t (21) = 6.760, Cohen’s d = 1.441; 0.5 s: t (18) = 8.883, Cohen’s d = 2.038; 1 s: t (20) = 7.199, Cohen’s d = 1.571; 2 s: t (21) = 5.010, Cohen’s d = 1.068, all p-values <.001] with no difference between ITIs [F(4,72) = 0.141, p-value = 0.966, ⍵² = 0]. An inverse relationship between previous trial time and temporal error variability - evidenced by β_2_ coefficients - was also found, but only for the 2 s ITI [0 s: t (21) = -1.380, Cohen’s d = -0.294, p-value = .182; 0.25 s: t (21) = -1.265, Cohen’s d = -0.270, p-value = .220; 0.5 s: t (18) = -0.853, Cohen’s d = -0.196, p-value = .405; 1 s: t (20) = -2.004, Cohen’s d = -0.437, p-value = .059; 2 s: t (21) = -2.887, Cohen’s d = -0.613, p-value = .009]. For the longest inter-trial interval, a longer time to contact in the previous trial led to a smaller variation in the response emitted in the current trial. No difference was found between ITIs regarding previous trial regressors for response variability [F(4,72) = 0.586, p-value = .674, ⍵² = 0].

Lastly, we compared the temporal error between the previous laboratory (-64.73±0.43, mean±SEM) experiment (Bilacchi et al., 2022) and the online (27.86±0.54, mean±SEM) experiment. Overall, temporal errors were more delayed [online - laboratory, t(43) = 4.63, p-value < .001] and more variable [online - laboratory, t(43) = 3.49, p-value = .001] in the online replication than in the laboratory experiment (figure 4). We then pooled together the serial dependence values calculated from Bilacchi et al. (2022) and the present data to fit a linear mixed model. We found a significant main effect of ITI [F(7,167.5) = 2.659, p-value = .012] but not of experiment type [F(1,74.2) = 0.333, p-value = .565], indicating both modulation of serial dependence strength between 0 and 8 s of ITI and no difference in serial dependence estimates from laboratory versus online produced data. A Bonferroni-Holm corrected post-hoc test showed a significant difference only between ITIs of 0.1 and 8 s [ITI_0.1_ - ITI_8_ = 0.07366, t(169) = 3.68481, p-value = .009], revealing a significantly smaller serial dependence after an 8 s ITI compared to blocks with an ITI of 0.1 s.

**Figure 4.**
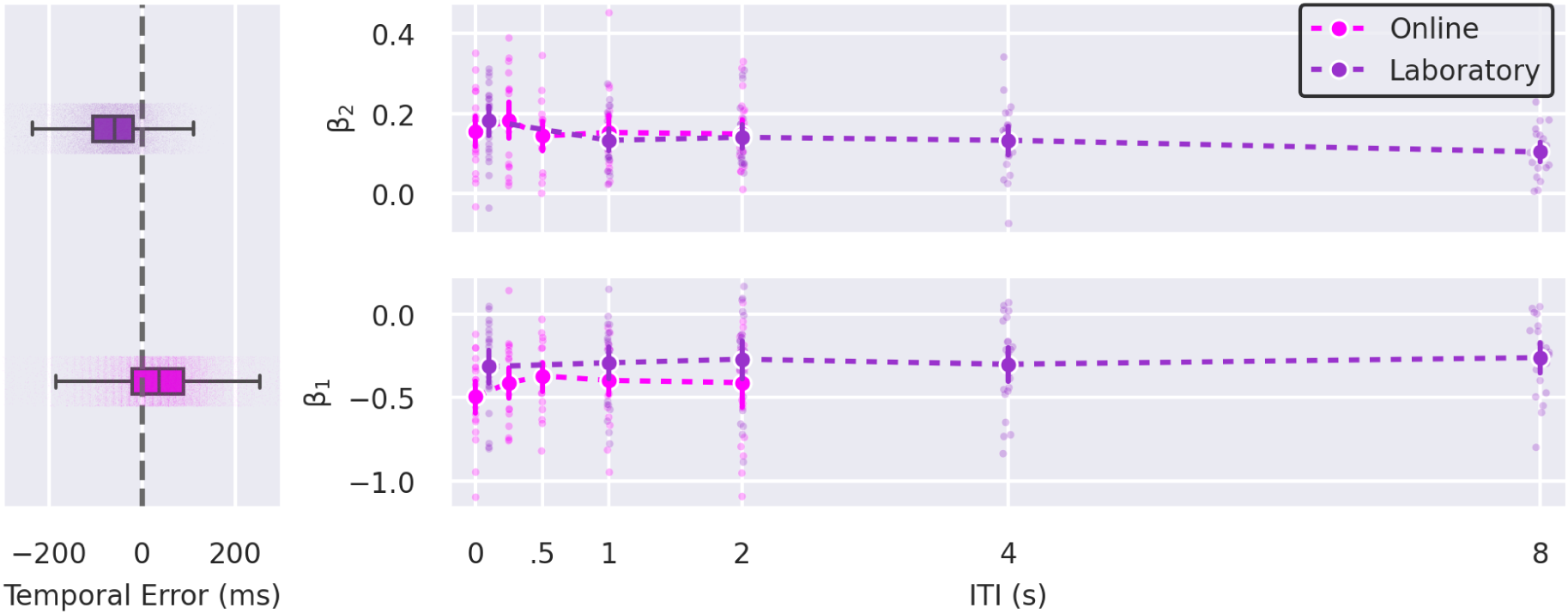
Data Comparison Between Laboratory and Online Data. *Note.* Temporal errors are more delayed and variable in online settings compared to laboratory (left). Current (right, bottom) and, more importantly, previous (right, top) trial biases are comparable between online and laboratory settings.

### Partial discussion

The results of Experiment 1 partially replicate and extend our previous findings (Bilacchi et al., 2022). As in the earlier study, we observed significant serial dependence across all ITI conditions. However, contrary to our previous results, we did not find significant modulation of serial dependence as a function of the ITI using an ANOVA (see below for a discussion of the combined data). Upon further analysis with a Helmert contrast, we found that serial dependence decreased after a 0.25 s ITI, in line with the previously observed decrease between 0.1 and 1 s ITIs (Bilacchi et al., 2022). However, this result should be interpreted with caution, as it was exploratory and did not follow from a significant main effect of ITI.

We also found a significant current trial bias for all ITI conditions, with a tendency for shorter responses for longer current trial intervals, as in Bilacchi et al. (2022). However, contrary to these previous results, the magnitude of this bias differed between ITIs due to a larger bias in the condition without an interval between trials. A larger current trial bias, irrespective of past information following a non-stop stream of trials, may be due to attention and the reduced time needed to prepare for the following response.

Regarding the variability of responses, we replicated the finding that longer target times lead to increased response variability in the current trial, with no differences across ITIs. In Bilacchi et al. (2022), we found that longer target times in the previous trial led to a decreased response variability across all ITIs. In the present study, however, we replicated this finding only for the 2 s ITI condition, likely due to already more variable responses being acquired in an online data collection modality.

Lastly, we employed a linear mixed model to combine results from our previous laboratory experiment (Bilacchi et al., 2022) and the current online experiment. Although responses are more variable in the online experiment, we did not observe differences in the size of serial dependence across tasks. When both experiments were analyzed together, we observed a modulation of serial dependence across ITIs. This difference was limited to comparing the ITI with the strongest serial dependence (in ITI 0.1 s) with the ITI with the weakest serial dependence (in the ITI of 8 s). Although this is an interesting finding, as it could indicate that serial dependence is strongest after very short ITIs, our experimental design and analysis were not planned to examine such comparisons. Future studies can systematically change the ITIs within more fine-grained values and use proper modeling that considers the fact that ITIs are continuous to study a possible modulation of the effect of the ITIs.

Taken together, our results indicate that serial dependence is robust to the passage of time, can be captured using an online setting of the visuomotor task employed in our previous studies (Bilacchi et al., 2022), and can last at least 8 seconds with a small decrease at most. In experiment 2, we alternated temporal coincidence trials with different interfering tasks to investigate further how robust serial dependence is and whether its strength depends on specific task demands.

## Experiment 2

### Materials and Methods

#### Participants

Twenty-eight participants (16 women, 26.607 ± 6.618 years old, mean ± standard deviation) were informed about the experimental procedures and agreed to a Consent Form provided via electronic correspondence. We recruited a convenience sample through advertising on social media. The experimental protocol was approved by The Research Ethics Committee of the Federal University of ABC. All experiments were performed following the approved guidelines and regulations. All participants had normal or corrected to normal vision and reported no acute neuropsychological disorders. The sample size was estimated using the same power analysis as in Experiment 1. Additional participants were recruited to compensate for exclusions when these were identified in the earlier phases of the preprocessing pipeline.

#### Stimulus and Task

Participants performed a similar coincident timing task as in Experiment 1, in which they pressed a button at the same time a target moving from left to right at a constant speed hit a barrier (figure 1). This was done in three sessions; in each session, a different task was performed between the coincident timing trials. The experiment was performed online, so screen size and room conditions varied. However, participants were instructed to be in a quiet environment. The experiment was built in *Psychopy* (v 2020.1.2; Peirce et al., 2019) and ran online through the *Pavlovia* platform.

In every session, we employed a calibration routine to assess participants’ screen size by requiring them to resize an image to match the dimensions of a physical credit card (Morys-Carter, 2022). In coincident timing trials, the target was a red square with a side of 0.59 cm and was initially drawn at the left, 0.42 cm above the vertical center of the screen. A semi-square the same size as the target marked the starting zone. The target was shown for 200 ms, marking the start of the trial, and then reappeared 200 ms later, starting a trajectory towards a vertical barrier of 30% the height of the screen on the right side. The target always moved at a constant speed, with interception times of 496, 450, 412, 379, or 351 ms. The target path was centered on the horizontal center of the screen, and participants were instructed to look at a fixation cross of the same size as the target during the whole trial and press the spacebar at the same time the target hit the barrier. Responses were recorded for up to 500 ms after the target disappeared behind the barrier and were followed by an inter-trial interval of 1 s.

Intervening tasks could be *Time Reproduction*, *Speed Judgment,* or *Orientation Judgment* (see the complete scheme in figure 1). Participants were instructed to answer the intervening tasks using their left hand over their keyboard’s Q and W keys and to maintain their right hand over the space bar to emit responses in the main task.

The *Time Reproduction* task consisted of three red squares of 0.59 cm that appeared on the left, at the top, and on the right of the fixation cross at a distance of 0.59 cm. Every square appeared for 200 ms. The left square appeared first, followed by the top square with an inter-stimulus interval of 496 ms or 351 ms (longest and shortest intervals of the main task). Participants were required to press the Q key on the keyboard to produce the right square after the same time had elapsed.

The *Speed Judgment* task consisted of a reference and a probe stimulus, both comprising a train of targets with a size similar to the one in the interception task (red square of 0.59 cm side), moving at a constant speed and continuously disappearing behind the barrier and reappearing at the start position. The reference speed was the same as the central speed for the interception task, i.e., each individual target of the target series took 412 ms to complete the trajectory. The probe speed could assume any of the speeds used in the interception task and always appeared after the reference. Each speed sample was shown for 500 ms, separated by 500 ms each. Participants were then instructed to answer via keypress with Q if the reference was faster than the probe and W if the probe was faster than the reference.

The *Orientation Judgment* task consisted of two successive bars (length 11.884 cm) that appeared for 200 ms, separated by a 600 ms noise mask. The first bar had a random orientation between 0° and 180°, whereas the second bar could be tilted 0°, 2.5°, 5°, 7.5°, 10°, 12.5° or 15° to either side. Participants were required to answer by pressing the Q key of the keyboard if the second bar was more counter-clockwise than the first or the W key if it was more clockwise.

#### Analysis

Temporal errors were calculated similarly to those in Experiment 1, and the same procedure was used to mark trials as outliers (Leys et al., 2013; Rousseeuw & Croux, 1993). One participant was excluded from further analysis for having more than 15% of the main task trials missed or marked as outliers, and another for exhibiting a median temporal error above 200 ms, which might indicate a poor task execution - a reaction to the moment of interception. Additionally, a linear (in the time reproduction task) or a binomial (for both the judgment tasks) regression was adjusted to the concurrent task data to verify whether participants performed the task correctly. Three additional participants were excluded with this procedure due to poor performance in at least one of the intervening tasks.

As in Experiment 1, serial dependence was accessed by modeling the TE for each participant and condition using a multiple linear regression (equation 1) in which β_2_ measures the influence of the previous trial on the current TE. Serial dependence in each condition was evaluated with a one-tailed, one-sample t-test on the β_2_ coefficients. We used repeated measures ANOVA to evaluate if different tasks modulated serial dependence. Biases associated with the current trial were assessed using a two-tailed t-test on the β_1_ coefficients. We performed data pre-processing and regressions in Python and statistics analysis in JASP (v 0.17.2).

### Results

Participants’ adjusted slopes from the intervening task were tested against zero with a one-sample t-test yielding significant results. These results indicate that participants perceive and respond to the target interval differences in the timing reproduction task and to the perceptual differences in speed and orientation in the judgment tasks [*Time*, t(22) = 6.928, Cohen’s d = 1.114; *Speed*, t(22) = 14.289, Cohen’s d = 2.043; *Orientation,* t(22) = 10.573, Cohen’s d = 1.114; all p-values < 0.001] (figure 5D).

**Figure 5.**
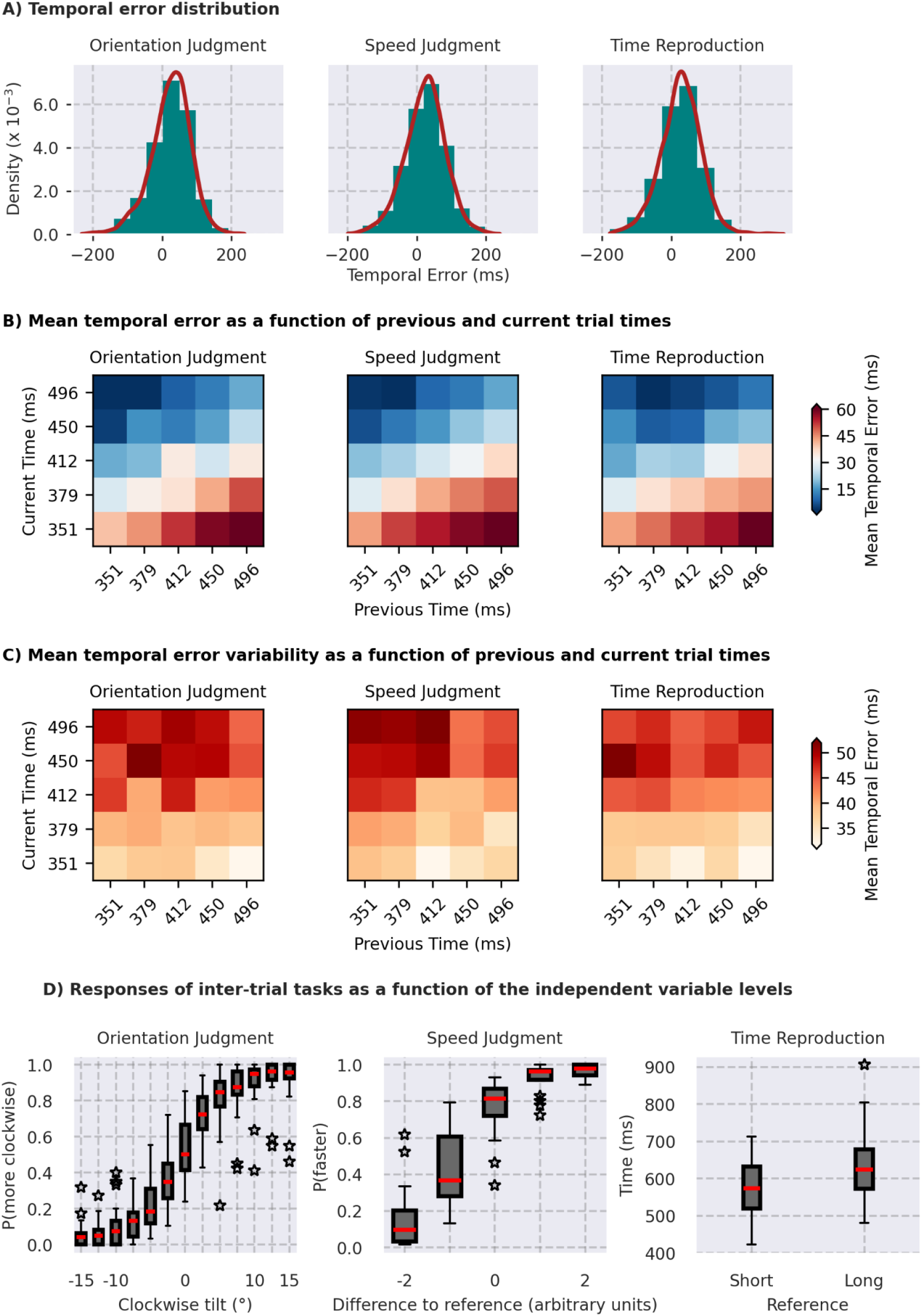
Data Overview of Experiment 2. *Note.* **A** temporal error density for all participants per inter-task condition. The distribution around 0 and between 200 and -200 milliseconds shows participants are performing the task as expected. **B** Mean temporal error for all volunteers as a function of previous (horizontal axis) and current (vertical axis) target times. Participants tend to anticipate targets that take longer to hit the barrier (cells get bluer from bottom to top), but also tend to delay answers as the previous target time increases (cells get redder from left to right). **C** Mean temporal error variability for all volunteers as a function of previous (horizontal axis) and current (vertical axis) target times. Responses become more variable as current target times increase (cells get redder from bottom to top), consistent with the scalar property of time. **D** Responses for the inter-trial tasks. Orientation judgments (left) have a higher probability of being ‘clockwise’ as the targets are tilted more clockwise. Speed judgments (center) have a higher probability of being ‘faster’ as the probe becomes faster than the reference. Time reproductions (right) are longer for the long reference intervals than for the short reference.

Participants’ responses were centered around 0 ms for all conditions (figure 5A), indicating they were neither anticipating nor reacting to the target. For each pair of current and previous times, we plotted TE’s means and standard deviations (figures 5B and 5C, respectively). Figure 5B exhibits the same general patterns seen in experiment 1: (1) columns become bluer from bottom to top as participants anticipate responses for longer times in the current trial, and (2) lines become redder from left to right as participants tend to answer later following longer times in the previous trial. As for the answers’ variability (figure 5C), no distinguishable pattern other than an increased response variability for longer times can be readily seen. Again, we inspected these patterns with a multiple linear regression (equation 1) fitted separately for each participant and condition.

For the coincident timing task, we found attractive serial dependencies in all conditions [*Speed*: t(22) = 9.396, Cohen’s d = 1.959; *Time*: t(22) = 4.806, Cohen’s d = 1.002; *Orientation*: t(22) = 9.834, Cohen’s d = 2.051, all p-values < 0.001] (figure 6), suggesting that the temporal errors for the current trial became more positive as time to contact increases in the previous trial independently of the task participants are executing between trials. The magnitude of this dependence was not modulated by the interfering task, as verified by a repeated measures ANOVA [F(2,44) = 1.766, p = .183, ⍵² = 0.017]. These results suggest that the intervening tasks employed could neither abolish nor modulate serial dependence.

**Figure 6.**
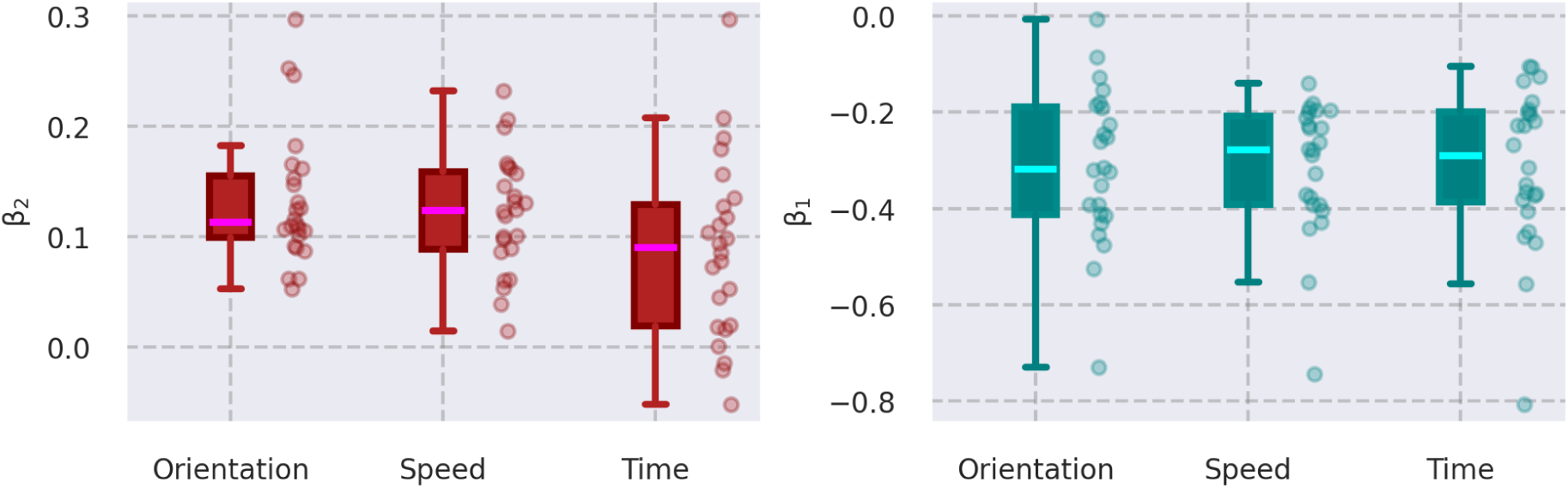
Linear Regression Slopes for Experiment 2. *Note.* Previous (left) and current (right) time slopes of multiple linear regressions performed on individual participant data for blocks of each inter-trial task. The further slopes are from zero, the bigger the contribution of the previous or current times for temporal error. Small transparent dots represent individual data.

Similarly, we tested if current trial biases generally found in coincident timing tasks were present, with significant results [*Speed*: t(22) = -8.682, Cohen’s d = -1.810; *Time*: t(22) = -7.018, Cohen’s d = -1.463; *Orientation*: t(22) = -6.596, Cohen’s d = -1.375, all p-values < 0.001] (figure 6), indicating that participants tend to respond earlier for longer target times on the current trial. Additionally, we found no difference in current trial biases between tasks [F(2,44) = 0.152, p = 0.859, ⍵² = 0].

Even though serial dependence remains unaltered, response variability could change, reflecting more or less demanding intertrial tasks. First, we compared the standard deviation of temporal errors pooled from each participant separately for each condition, with no detectable difference [F(2,44) = 1.927, p = .158, ⍵² = 0.008]. Then, as in experiment 1, we calculated the standard deviation from temporal errors per participant for each pair of current and previous times and fitted a similar multiple linear regression as in equation 1. The variability of responses increased with current time to contacts in all conditions [*Speed*: t(22) = 6.841, Cohen’s d = 1.426; *Time*: t(22) = 6.055, Cohen’s d = 1.263; *Orientation*: t(22) = 6.469, Cohen’s d = 1.349, all p-values < 0.001] (see pattern in figure 5C), and this relationship was not different across conditions [F(2,44) = 1.614, p = .211, ⍵² = 0.009]. Similarly, response’s variability decreased with previous times to contact [*Speed*: t(22) = -3.646, Cohen’s d = -0.760, p-value = .001; *Time*: t(22) = -2.663, Cohen’s d = -0.555, p-value = .014; *Orientation*: t(22) = -2.354, Cohen’s d = -0.491, p-value = .028], but once more there was no difference in this decrease between conditions [F(2,44) = 1.299, p = .283, ⍵² = 0.009]. That is, the proportionality between temporal error variability and time to contact in the current trial observed in experiment 1 is maintained even with concurrent tasks, whilst variability decreases with previous trial time in all conditions in experiment 2, and the nature of these tasks does not change this relationship.

Lastly, we regressed temporal errors per participant and session again - as in equation 1 -, this time adding information from the concurrent task of the last trial. For the judgment tasks (speed or orientation) this regressor was the difference between the reference and the probe in degrees for the orientation task (15, 12.5, 10, 7.5, 5, 2.5 or 0 degrees clockwise or counter-clockwise) and arbitrary units for the speed task (-2, -1, 0, 1 or 2). For the timing reproduction task, this regressor was the target interval in the last trial (either 496 or 351 ms). We did not find the regressors for intertrial tasks to be different from zero for the speed and judgment tasks [*Speed*: t(22) = -0.262, Cohen’s d = -0.055, p-value = .796; *Orientation*: t(22) = 1.431, Cohen’s d = 0.298, p-value = .166], but we found that the temporal error on the current trial becomes more positive after participants are required to reproduce the longer reference time in the time reproduction task [t(22) = 2.577, Cohen’s d = 0.537, p-value = .017]. These results show that responses were impacted by previous time reproductions in the intervening task, even though the presence of this task did not abolish or weaken serial dependence.

### Partial discussion

In Experiment 2, the goal was to selectively disrupt serial dependence to understand the nature of the memory traces leading to the bias and further investigate the effects of the intervening tasks on visuomotor integration. Firstly, we found serial dependence in all conditions with no differences, showing that the bias during visuomotor integration is robust to interference from both motor and perceptual tasks. When adding information from concurrent tasks to the regression used to estimate serial dependence, we found an influence only for the temporal and not for the judgment tasks, suggesting that the nature of the information being reproduced - a time duration - is important for this interference to occur. We also found current trial biases in this experiment, with longer trials leading to more premature responses. These results confirmed the findings regarding current trial biases in experiment 1 and previous experiments (Bilacchi et al., 2022), and it was not different across conditions.

On the same line of thought that memory traces in experiment 1 might be weakened in their precision but not in their magnitude, we also studied response variability in experiment 2. Overall response variability was not different between conditions. We also found that response variability increases as a function of the current trial time, possibly due to the scalar property of time. Contrary to experiment 1 and in accordance with previous findings (Bilacchi et al., 2022), we found an inverse relationship between previous trial time and current response variability in all conditions.

## Discussion

Our primary goal was to investigate the nature of memory traces underlying serial dependence in a visuomotor integration task involving coincident timing. First, we aimed to replicate and expand upon previous findings regarding the decay of these memory traces. Second, we sought to determine the type of information retained between trials by introducing intervening tasks. These tasks were designed to potentially disrupt serial dependence if they recruited the same neural resources required to store the memories from previous trials.

The duration of serial dependence has been examined in previous studies, with some finding effects lasting up to four prior trials, or approximately 15 seconds (J. Fischer & Whitney, 2014; Kalm & Norris, 2018). However, when intervening trials were removed to examine serial dependence dynamics, mixed results were found. Bliss et al. (2017) observed a sharp decrease in serial dependence between 3 and 6 seconds, whereas Papadimitriou et al. (2015) reported more stable serial dependence within the same time frame for macaques. It is important to note that all these studies used delayed memory tasks that required explicit stimulus feature encoding during the delay period.

For coincident timing tasks, our group tested a range of inter-trial intervals and observed the maintenance of serial dependence for up to 8 s after an abrupt decrease between 0.1 and 1 s (Bilacchi et al., 2022). Previously, we raised the possibility of a change in storage mode for the memory responsible for serial dependence, resulting in this sudden change, followed by a stable bias. In our current study, in experiment 1, we observed that this change occurred between 0.25 and 0.5 s, although this decay was less pronounced than in our previous study. This profile is more congruent with the time constant of a decay of uncertainty associated with memory traces, an alternative hypothesis we have also previously suggested (Bilacchi et al., 2022).

Working memory can keep the uncertainty associated with the memorized stimulus (Honig et al., 2020), which itself has a decaying profile that could affect serial dependence. Following Bayesian principles (Van Bergen & Jehee, 2019), the influence of past trials on the current trial should be maximized when the previous trial uncertainty is low and the current trial uncertainty is high. However, evidence only supports the latter, while the former seems to play no role (Ceylan et al., 2021; Cicchini et al., 2018). This could be attributed to a rapid decay of uncertainty associated with memoranda to the point that it does not affect serial dependence in delayed memory task paradigms used in these works. In our task, however, the time between the stimulus presentation and the next trial comprises only the inter-trial interval, which was virtually instantaneous in one of our conditions. Perhaps, by getting rid of retention periods and having the stimulus on-screen during the whole trial, our task was able to capture the effects of encoded uncertainty of the last trial during shorter ITIs. As the information uncertainty for the last trial fades, serial dependence becomes stable after 1 s. This, however, would need further studies to be asserted, as the uncertainty of the previous trial was not manipulated here.

Serial dependence is influenced by contextual information (Feigin et al., 2021), stimulus location (J. Fischer & Whitney, 2014), attention to stimuli (J. Fischer & Whitney, 2014; Fritsche et al., 2017), and stimulus selection during visual search (Rafiei et al., 2021). These studies suggest that serial dependence plays a role in binding stimuli features across space and time (Manassi & Whitney, 2024), with attended targets showing attractive serial dependence and non-target stimuli exerting either no influence or a slight repulsive bias. For instance, Houborg et al. (2023) found that participants instructed to ignore an intervening task’s orientation stimulus showed a repulsive bias on the main orientation reproduction task. In contrast, our experiment 2 showed no evidence that intervening tasks disrupt serial dependence and instead found a small attractive bias caused by the stimuli in the temporal judgment task. We will discuss possible explanations for these results in the following paragraphs.

The intervening trials in our second experiment involved stimuli that were physically similar to the main task’s target. The Time Reproduction condition had a temporal interval similar to the main task’s time-to-contact, and the Speed Judgement condition used the same target speeds as the main task. While recent research suggests that physically similar stimuli across trials create a repulsive effect (Moon & Kwon, 2022; Pascucci et al., 2019; Sadil et al., 2024; Sheehan & Serences, 2022), especially when stimuli need to be ignored (Houborg, Pascucci, et al., 2023; Rafiei et al., 2021), our second experiment showed no significant interference in the main task. This suggests that physical similarity alone doesn’t cause a repulsive bias. Unlike studies where participants actively ignored distractors, our participants interacted with and used stimulus information to perform the intervening task. This difference may explain the absence of a repulsive bias influencing responses. Additionally, the lack of bias in the Speed Judgement task might be due to the canceling out of repulsive and attractive biases. Our results emphasize that bias formation is complex and depends on task interaction, not just stimulus characteristics.

Researchers have long proposed that sensory systems contribute to memory storage (Christophel et al., 2017; Fuster, 1997). If the same neural resources are involved in processing incoming information and maintaining it across trials, intervening tasks should introduce interference and disrupt serial bias. However, in Experiment 2, we observed no reduction in serial dependence across different intervening tasks. This finding suggests that incoming stimuli do not compete for the same neural resources used to retain memory traces from the previous trial. For instance, processing speed in the Speed Judgment task may rely on distinct neural populations from those supporting the short-term memory storage of speed in the main task. This result challenges the core premise of the sensory recruitment hypothesis, which posits that the same sensory regions responsible for processing incoming stimuli also store them in memory. While sensory areas may still contribute to memory storage, they likely do so via a separate neural population or an alternative coding scheme (Adam et al., 2022).

One possible explanation is that both memories are stored within the same neural populations at different moments. Evidence suggests that visual areas can simultaneously process new distractor information while maintaining short-term memory traces (Rademaker et al., 2019). Additionally, research indicates that information can be silently stored in synaptic weights (Mongillo et al., 2008; Rose et al., 2016; Stokes, 2015; Wolff et al., 2015, 2017) and later reactivated into firing patterns when needed (Barbosa et al., 2020). Theoretically, this dynamic interplay between activity-silent and active states could protect stored information from interference. However, this remains speculative and could be tested using electroencephalogram recordings during task performance. Conversely, the serial bias observed in the Temporal Reproduction condition suggests that, at least for time perception, processing new information and maintaining previous trial data may rely on overlapping neural resources. Similarly, Bueno et al. (2024) found that temporal judgment and reproduction tasks engage similar neural mechanisms. However, this interaction does not appear disruptive; serial dependence persists without significant weakening or erasure.

Motor components recruited for response emission during the time reproduction task may interfere with response on the subsequent trial regardless of perceptual representations of the moving target in the main task. That is, two different neural populations are dealing with the information from each task, but both interfere with responses. Serial dependence was initially described as a low-level perceptual phenomenon, occurring even without responses on the last trial (J. Fischer & Whitney, 2014). This view is not undisputed, as recent evidence suggests that the bias arises from decisional stages (Ceylan et al., 2021; Feigin et al., 2021; Fritsche et al., 2017) and then propagates to early levels of sensory processing (Cicchini et al., 2021). In the case of the task used in the present study, interference with the premotor cortex weakens - but does not extinguish - serial dependence (De Azevedo Neto & Bartels, 2021). Recent models suggest that serial dependence results from both an attraction toward prior responses and a subtle repulsion from prior stimuli (Moon & Kwon, 2022; Pascucci et al., 2019; Sadil et al., 2024; Sheehan & Serences, 2022), although behavioral and electrophysiological data indicate a repulsive effect of prior responses (Zhang & Luo, 2023). However, a purely motor explanation for our results is unlikely, as categorical responses in the Orientation and Speed Judgment tasks would also influence temporal errors in the main task, which our analysis does not capture. Instead, our results align with the idea that history biases arise from multiple influences (Manassi & Whitney, 2024), given that temporal reproductions partially explain the variability of temporal errors without affecting serial biases. Notably, our time reproduction task included only two levels of the independent variable to limit experiment complexity and duration. A more detailed investigation is needed to assess how time reproductions interfere with serial dependence during consecutive target interceptions.

During target interception, multiple stimulus features — such as target speed, distance traveled, and time duration before reaching the barrier — provide information for response timing. This redundancy may help preserve serial dependence despite interference. Since serial biases can occur independently for different stimulus features (Zhang & Luo, 2023), their overlap may mitigate disruptions. Future studies should explore how simultaneous stimulus features shape serial dependence in visuomotor integration tasks. Lastly, since the intervening and main tasks were entirely different, there might have not been any coupling or sense of continuity between stimuli (Manassi & Whitney, 2024), preventing memory traces from merging with incoming sensory information. Furthermore, recent evidence suggests that serial dependence is more likely when the processed information is relevant to current task goals (C. Fischer et al., 2024; Houborg, Kristjánsson, et al., 2023), which may explain why the intervening tasks had little impact on the main task.

Our results show that stable, long-lasting memory traces drive serial dependence during visuomotor integration, remaining robust to interference. None of our manipulations abolished or diminished serial dependence. Even performing an unrelated task with entirely different stimuli (e.g., orientation) does not reduce serial dependence, demonstrating the effect’s robustness to task-switching and distraction. Intervening temporal reproductions, however, induced serial biases in the main task. We speculate that the stimuli’s redundant multi-feature nature enhances its resistance to interference. Future studies should examine whether the timing or motor demands of the intervening task influence responses. Manipulating the difficulty of intervening timing tasks or the complexity of responses may help identify conditions under which serial dependence can be disrupted. Additionally, researchers can explore delayed memory paradigms with redundant multi-feature stimuli that guide behavior to clarify how information redundancy supports the maintenance of serial biases. Understanding these mechanisms will provide deeper insights into the stability of serial dependence and its implications for visuomotor behavior.

## Declarations

### Funding

E.V.P.S. was supported by grant #2019/06423-7, São Paulo Research Foundation (FAPESP). A.M.C. was supported by Grant #2017/25161–8, São Paulo Research Foundation (FAPESP) and by grants #311721/2023-0 and #407769/2023-4, Conselho Nacional de Desenvolvimento Científico e Tecnológico (CNPq), Brasil.

### Conflicts of interest

The authors have no competing interests to declare that are relevant to the content of this article.

### Ethics approval

Approval was obtained from the Research Ethics Committee of UFABC (CAAE: 84603118.8.0000.5594).

### Consent to participate

All participants provided informed consent through a Consent Form provided via electronic correspondence. The Consent Form was approved and overseen by the Research Ethics Committee of UFABC.

### Consent for publication

Not applicable.

### Availability of data and materials

The datasets generated during the current study, materials and analysis scripts are available in the Open Science Framework repository, https://osf.io/2j4ze/.

### Code availability

Not applicable.

## Notes

https://osf.io/2j4ze/?view_only=b939898b7e9144c3aa46f4f85020c51b

### Competing Interest Statement

The authors have declared no competing interest.

